# STAT3 cooperates with the core transcriptional regulatory circuitry to drive *MYC* expression and oncogenesis in anaplastic large cell lymphoma

**DOI:** 10.1101/2022.08.31.506044

**Authors:** Nicole Prutsch, Shuning He, Alla Berezovskaya, Adam D. Durbin, Neekesh V. Dharia, Kimberly Stegmaier, Jamie D. Matthews, Lucy Hare, Suzanne D. Turner, Lukas Kenner, Olaf Merkel, Richard A. Young, Brian J. Abraham, A. Thomas Look, Mark W. Zimmerman

## Abstract

Anaplastic large cell lymphoma (ALCL) is an aggressive, CD30^+^ T-cell lymphoma of children and adults. *ALK* fusion transcripts or mutations in the JAK-STAT pathway are observed in most ALCL tumors, but the mechanisms underlying tumorigenesis are not fully understood. Here we show that dysregulated STAT3, together with a core transcriptional regulatory circuit consisting of BATF3–IRF4– IKZF1, co-occupies gene enhancers to establish an oncogenic transcription program and maintain the malignant state of ALCL. Critical downstream targets of this network in ALCL cells include the proto-oncogene *MYC*, which requires active STAT3 to facilitate high levels of *MYC* transcription. The activity of this auto-regulatory transcription loop is reinforced by MYC binding to the enhancer regions associated with *STAT3* and each of the core regulatory transcription factors. These findings provide new insights for understanding how dysregulated signaling pathways hijack cell-type-specific transcriptional machinery to drive tumorigenesis and create therapeutic vulnerabilities in genetically defined tumors.

## Introduction

Anaplastic large cell lymphoma (ALCL) is a CD30+ subtype of T-cell lymphoma with around half of patients exhibiting chromosomal rearrangements involving the *ALK* gene^1–6^. Treatment with ALK inhibitors can achieve impressive complete responses in ALCL cells that express the *NPM1-ALK* fusion oncogene and with PDGFR inhibitors for a subset of these patients^7^. However, long-term ALK inhibition can lead to relapse and the overall survival for ALK-negative patients remains poor^8–11^. ALK-negative ALCLs have been found to harbor activating mutations in *JAK1, TYK2, ROS1*, and *STAT3*, each of which result in dysregulation of the JAK-STAT signaling pathway^12,13^. Targeted therapies with tyrosine kinase inhibitors such as JAK1/2 and TYK2 inhibitors have shown efficacy in preclinical models of genetically defined ALK-negative ALCL subsets^7,13,14^. Otherwise, treatment strategies for ALK-negative patients are limited to chemotherapeutic regimens such as CHOP (cyclophosphamide, doxorubicin, vincristine, prednisone), which are usually ineffective at managing the disease^15^.

The JAK-STAT pathway is a highly conserved signaling cascade activated by receptor tyrosine kinases in response to binding a wide range of cytokines and growth factors^16^. JAK-STAT and other signaling pathways provide cells with a mechanism for relaying information from the extracellular microenvironment to the nucleus, where transcriptional effectors such as STAT3 regulate gene expression in a cell-type-specific manner^17^. However, the mechanisms explaining how STAT3 interacts with cell-type-specific enhancers and how dysregulation of these pathways results in ALCL tumorigenesis are poorly understood.

In this study, we show activated of STAT3, a terminal effector of the JAK-STAT signaling pathway, interacts with a set of key transcription factors that form an interconnected autoregulatory loop termed the transcriptional core regulatory circuit (CRC) governing the ALCL expression program^18^. This autoregulatory transcriptional loop consists of BATF3, IRF4, and IKZF1, and drives high levels of expression of each of these transcription factors due to positive feedback that each transcription factor exerts on super- or stretch-enhancers (SE) associated with each of these genes^19,20^. As we and others have shown, STAT3 is activated downstream of ALK or due to direct mutation or fusion of genes including JAK1, JAK2, TYK2, or ROS1^12,13,21^. Here we show that aberrantly activated STAT3 binds concomitantly with each of the CRC transcription factors in small regions of ∼1 kb of open chromatin within their SEs as part of the autoregulatory loop of transcription factor genes, mediating high levels of expression of *MYC* and each member of the CRC. These results provide new insights into the mechanisms by which oncogenic signaling pathways collaborate with CRC transcription factors to drive the oncogenic gene expression program in ALCL.

## Results

### Core regulatory transcription factors and STAT3 cooperate to drive the ALCL gene expression program

Based on our prior work examining chromatin structures in pediatric leukemias and solid tumors, we began experiments to identify the transcriptional core regulatory circuit that mediates the oncogenic cell state in ALCL^22,23^. We first performed enhancer profiling based on ChIP-seq for H3K27ac because genes encoding CRC transcription factors are generally regulated by large *cis*-regulatory elements called super- or stretch-enhancers (SE). SEs are stretches of adjacent enhancer elements that concentrate large amounts of the transcriptional machinery into biomolecular condensates capable of driving high levels of gene expression from target promoters^19,24^. A group of 38 genes encoding transcription factors were consistently associated with SEs across at least five of eight analyzed ALCL cell lines, which include both ALK translocated and ALK-independent ALCL subtypes (Fig. 1a). Based on these results, we focused on the three transcription factors with lineage-restricted expression patterns – BATF3, IRF4, and IKZF1 – whose enhancer regions have consistently high levels of H3K27ac modification ALCL cells. ChIP-seq coverage tracks demonstrated that each of these genes is associated with a region covered very highly by acetylation at H3K27, meeting the criteria for SE in all eight ALCL cell lines, one ALCL patient-derived xenograft, and one primary patient ALCL (Fig. 1b-d). Next, we examined the expression levels of these three transcription factors in over 1200 cancer cell lines included in the Cancer Cell Line Encyclopedia^25^. We found that BATF3, IRF4, and IKZF1 are selectively highly expressed in transcriptomic analyses from each of the ten ALCL cell lines included in this data set, as highlighted in red in Figure 1e-g. Thus, the very high level of expression of these three transcription factors in ALCL, regardless of whether individual cell lines have a translocated and activated *ALK* gene, led us to postulate that *BATF3, IRF4*, and *IKZF1* form an essential core regulatory circuit (CRC) that determines cell state in ALCL.

**Figure 1.**
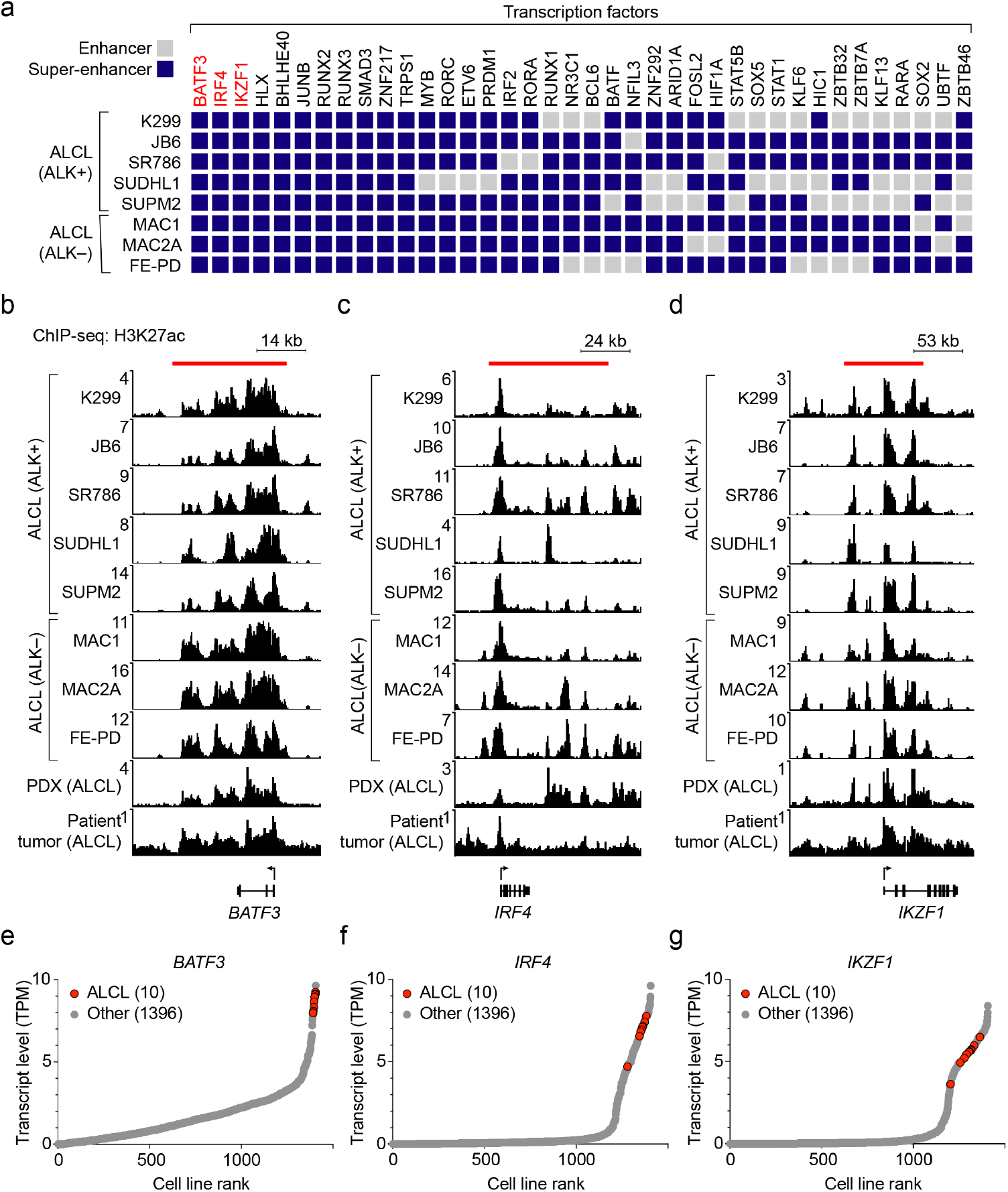
Highly expressed transcription factors are associated with SEs in human ALCL. a) Enhancer profiling of ALCL cell lines revealed a conserved set of SEs associated with transcription factor encoding genes (n=38). b-d) Normalized ChIP-seq alignment tracks for H3K27ac reveal SE regions associated with the *BATF3* (b), *IRF4* (c) and *IKZF1* (d) gene loci in ALCL. ChIP-seq read densities (y-axis) were normalized to reads per million reads sequenced from each sample. Red bars indicate the location of SEs. e-g) Transcript levels (y-axis) of *BATF3* (e), *IRF4* (f) and *IKZF1* (g), ranked by gene expression (x-axis) across cell lines. Ten ALCL cell lines (shown in red) are compared to n=1396 cell types derived from other lineages (shown in gray).

The CRC is an interconnected autoregulatory loop of SE-driven transcription factors that cooperatively regulate an extended network of genes that establishes cell state^18,26^. The enhancers associated with *BATF3, IRF4*, and *IKZF1* were each highly enriched in H3K27ac across all samples we examined, including NPM-ALK+ and ALK-negative cell lines (Fig. 2a). To test whether these transcription factors fulfill the interconnectivity requirements to qualify as CRC members, we performed CUT&RUN sequencing assays with antibodies specific for each of these factors to identify regions of sequence-specific genomic occupancy (Fig. 2b-d). For comparison, we also assayed the histone acetylation reader BRD4, and the MED1 component of the mediator complex. We found that BATF3, IRF4, and IKZF1 proteins each bind together within small regions of open chromatin called epicenters of ∼1 kb within their own and each other’s regulatory super-enhancers, establishing that they form an interconnected, autoregulatory transcriptional loop.

**Figure 2.**
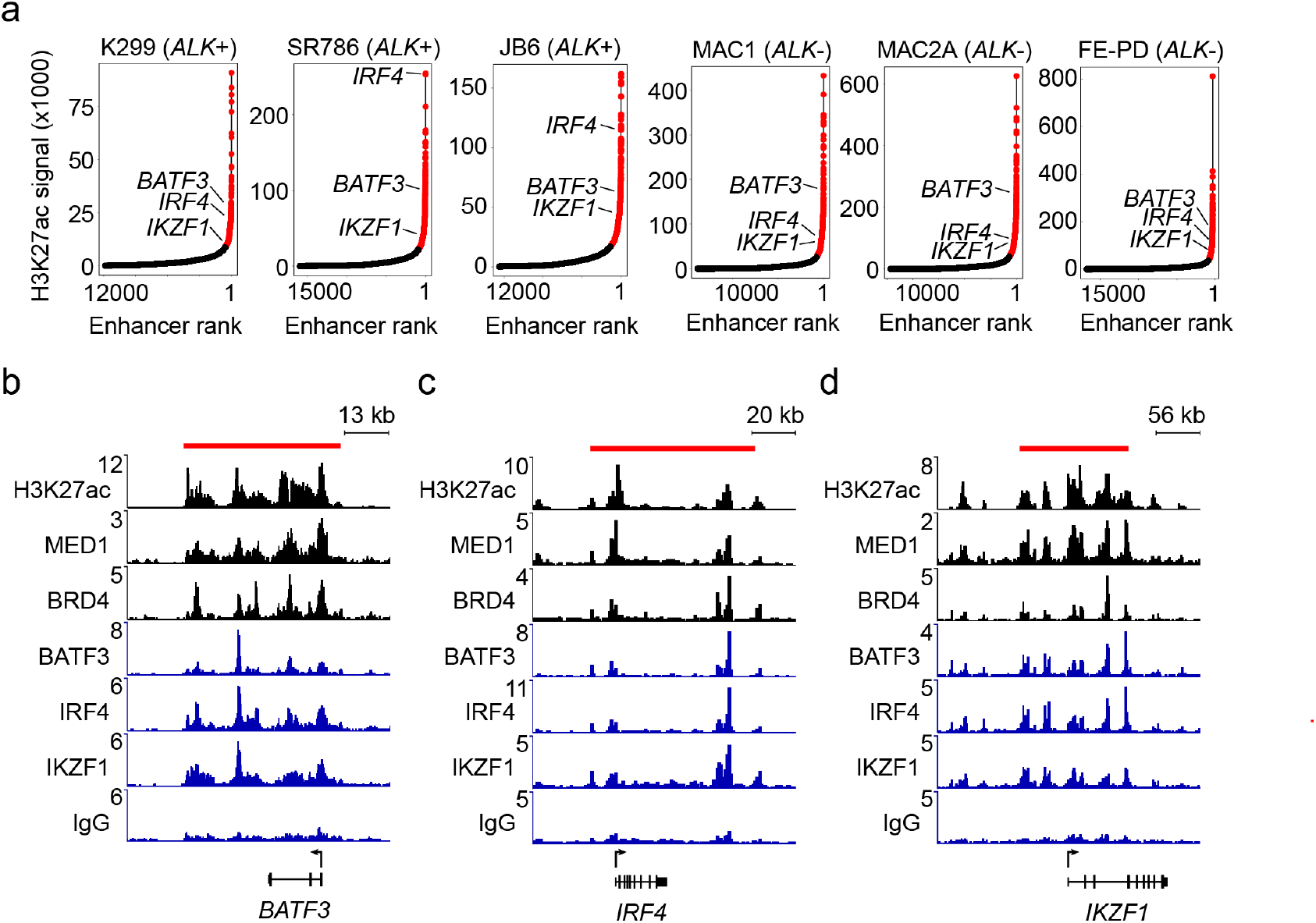
Identification and validation of an interconnected core transcriptional regulatory circuit in ALCL. a) Ranking of enhancers by H3K27ac signal associated with all genes in *NPM-ALK*+ (K299, SR786, and JB6) and *ALK*– (MAC1, MAC2A, and FE-PD) ALCL cells. *BATF3, IRF4*, and *IKZF1* were associated with super-enhancers across all ALCL cell lines. b-d) CUT&RUN sequencing alignment tracks for MED1, BRD4, BATF3, IRF4, IKZF1 and IgG (control) in MAC2A cells, overlaid with H3K27ac ChIP-seq, shown at the gene loci for *BATF3* (b), *IRF4* (c), and *IKZF1* (d). Read densities (y-axis) were normalized to reads per million reads sequenced in each sample.

### *STAT3* and the CRC transcription factors *BATF3, IRF4*, and *IKZF1* are selective gene dependencies in ALCL

Tumor-selective gene dependencies have been proposed as potential targets for cancer therapy because they are preferentially required for the growth and survival of particular tumor cells^27,28^. The DepMap Consortium has conducted a genome-scale CRISPR-Cas9 dependency screen in over 1000 cancer cell lines, including five ALCL cell lines^28,29^. We analyzed these results and found a small group of genes that each qualifies as a selective dependency in ALCL cell lines compared to cancer cells derived from other lineages. Notably, the core regulatory transcription factors *BATF3, IRF4*, and *IKZF1* were selectively essential in ALCL cells compared to cell lines derived from other tumor types (Fig. 3a). ALCL cells were also selectively dependent on the signaling transcription factor *STAT3*, a terminal effector of the JAK-STAT pathway that has recently emerged as an attractive target in ALCL (Fig. 3a)^12,30^. Two additional non-transcription factor selective gene dependencies included *PTPN2* and *SBNO2. PTPN2* encodes a protein tyrosine phosphatase known to dephosphorylate JAK1 and STAT3^31–33^. PTPN2 has also been reported to regulate ALK phosphorylation and activity, with a potential role in resistance to ALK inhibition^34^. Less is known about *SBNO2*, a transcriptional coregulator, which has been reported as a downstream target of IL-10 and STAT3 signaling in hematopoietic cells^35^. Overall, each of these selective gene dependencies in ALCL appears to be a member of the CRC or draws focus to the JAK-STAT signaling pathway as a crucial mediator of ALCL identity.

**Figure 3.**
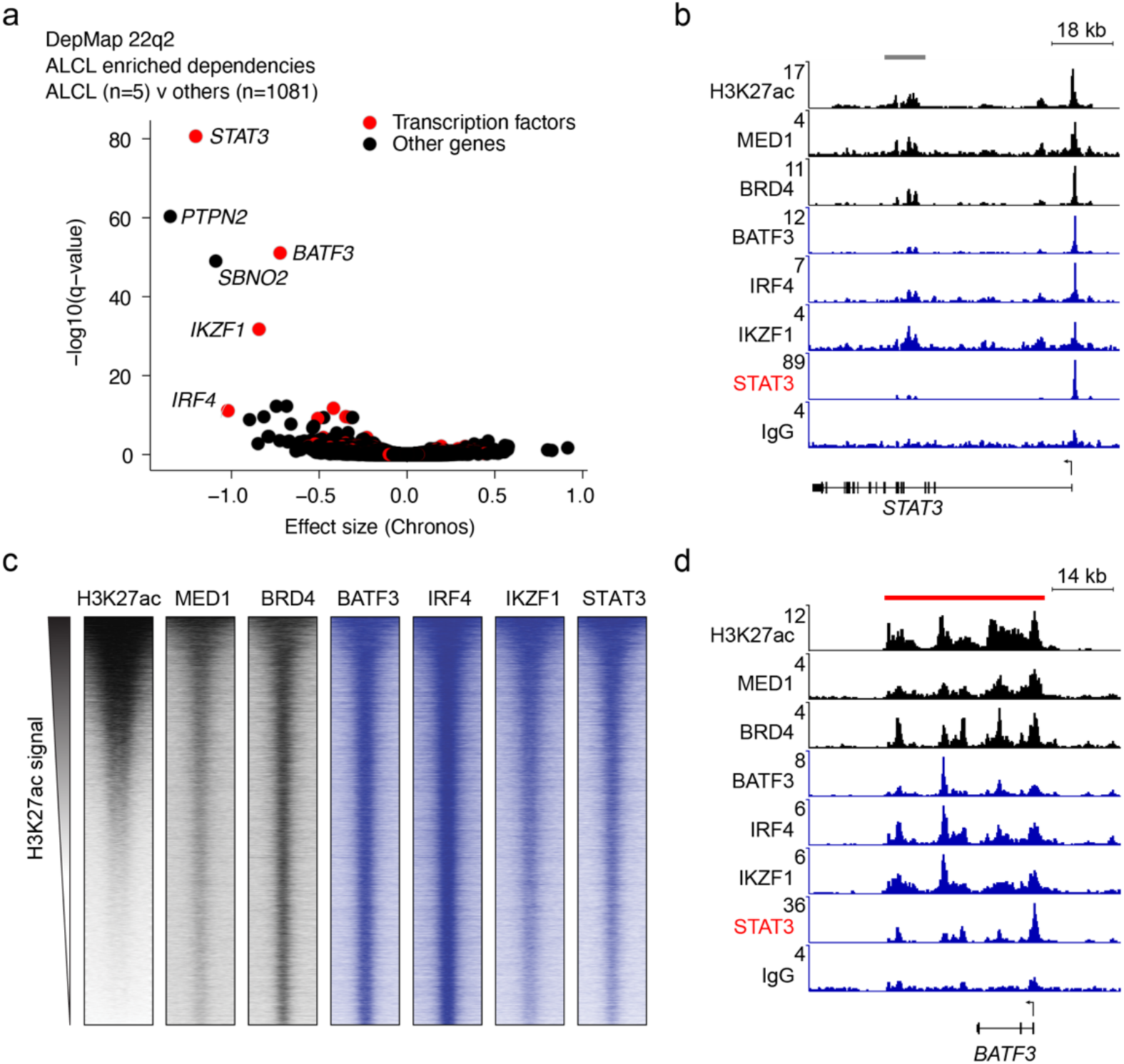
Tumor-selective gene dependencies in ALCL include the CRC and STAT3. a) Scatter plot showing dependency effect size (x-axis) and ALCL selectivity (y-axis) for all protein-coding genes. Genes encoding transcription factors are highlighted in red, and all other protein classes are shown in black. b) CUT&RUN sequencing alignment tracks for MED1, BRD4, BATF3, IRF4, IKZF1, STAT3 and IgG (control) in MAC2A cells, overlaid with H3K27ac ChIP-seq at the *STAT3* locus. Gray bar indicates the location of a typical enhancer peak. c) Genome-wide co-occupancy for H3K27ac, MED1, BRD4, BATF3, IRF4, IKZF1, and STAT3 determined by ChIP-seq/CUT&RUN sequencing. Genomic regions (rows) were defined as those enriched in sequencing reads for at least one target and are ranked by the H3K27ac signal therein. d) CUT&RUN sequencing alignment tracks for MED1, BRD4, BATF3, IRF4, IKZF1, STAT3 and IgG (control) in MAC2A cells, overlaid with H3K27ac ChIP-seq at the *BATF3* locus. Red bar indicates the location of a SE.

ALCL tumors frequently harbor driver translocations activating the *ALK* tyrosine kinase receptor or mutations activating components of JAK-STAT signaling pathways^12^. The terminal signaling effector of either event is the activation of STAT3, which becomes hyper-phosphorylated, dimerizes, and then enters the nucleus to bind to its genomic signal response elements to activate key target genes in a sequence-dependent manner^21,36–38^. Based on this information and our finding of selective *STAT3* dependency in ALCL, we focused on the role of STAT3 and its interaction with the CRC. We first examined the extent of localized H3K27ac by ChIP-seq, marking potential enhancers near the *STAT3* gene. The *cis*-regulatory enhancers associated with *STAT3* did not meet the quantitative H3K27ac enrichment threshold for a SE in most ALCL cells, despite high expression levels of both STAT3 transcript and protein in ALCL (Fig. 3b). Instead, high occupancy of BATF3, IRF4, IKZF1 and STAT3 was primarily detected at the *STAT3* promoter. Thus, *STAT3* lacks a conserved autoregulatory SE in ALCL cells and therefore did not meet our criteria for defining a CRC transcription factor gene.

Consistent with collaboration between STAT3 and the CRC, genome-wide occupancy analysis shows that the STAT3 protein binds concomitantly with BATF3, IRF4, and IKZF1 at H3K27ac modified regions throughout the genome (Fig. 3c). Unlike most transcription factors, STAT3 is dependent on post-transcriptional regulation through phosphorylation and dimerization downstream of activation of tyrosine kinases such as ALK or JAK family members^36^. Therefore, STAT3 functions as a signaling responsive transcription factor ordinarily dependent on the stimulation of receptor tyrosine kinases before it is activated to cooperate with CRC transcription factors in controlling gene expression^24^. This concept can also explain why the expression of *STAT3* is observed in many cell types, but dependency on STAT3 is restricted to ALCL and a limited number of other cell types where STAT3 signaling is active. In ALCL cells STAT3 occupancy was observed at SEs associated with CRC transcription factors indicating a role in reinforcing the expression of this positive feedback loop (Fig. 3d). These results demonstrate that dysregulation of STAT3 by NPM-ALK or other driver mutations allows it to function as a *de facto* CRC component in ALCL, collaborating with BATF3, IRF4 and IKZF1 to establish an oncogenic transcription program and malignant cell state.

### Activated STAT3 collaborates with CRC transcription factors to drive *MYC* expression

Upon activation, signaling transcription factors become highly concentrated at SEs that are co-occupied by core regulatory transcription factors^24^. In order to understand the effects of activated STAT3 on the tumor transcriptome, we performed spike-in normalized mRNA-seq analysis was performed on MAC2A (PCM1-JAK2+) cells treated with the JAK1/2 inhibitor ruxolitinib, and JB6 (NPM-ALK+) cells treated with the ALK inhibitor crizotinib, to determine fold-change in global transcript levels at 24 h (Fig. 4a,b). Notably, *MYC* transcript levels (highlighted in red) were among the most significantly downregulated in both cell lines. We performed ChIP-seq/CUT&RUN with an antibody specific for STAT3 in three ALCL cell lines – JB6 (ALK+), MAC2A (ALK-), FE-PD (ALK-), and an ALK+ PDX model – and found that STAT3 localizes with the CRC transcription factors BATF3, IRF4, IKZF1 at multiple epicenters across an SE region associated with the *MYC* gene (Fig. 4c,d). Thus, in ALCL STAT3 cooperates with all three CRC transcription factors in the regulation of MYC expression.

**Figure 4.**
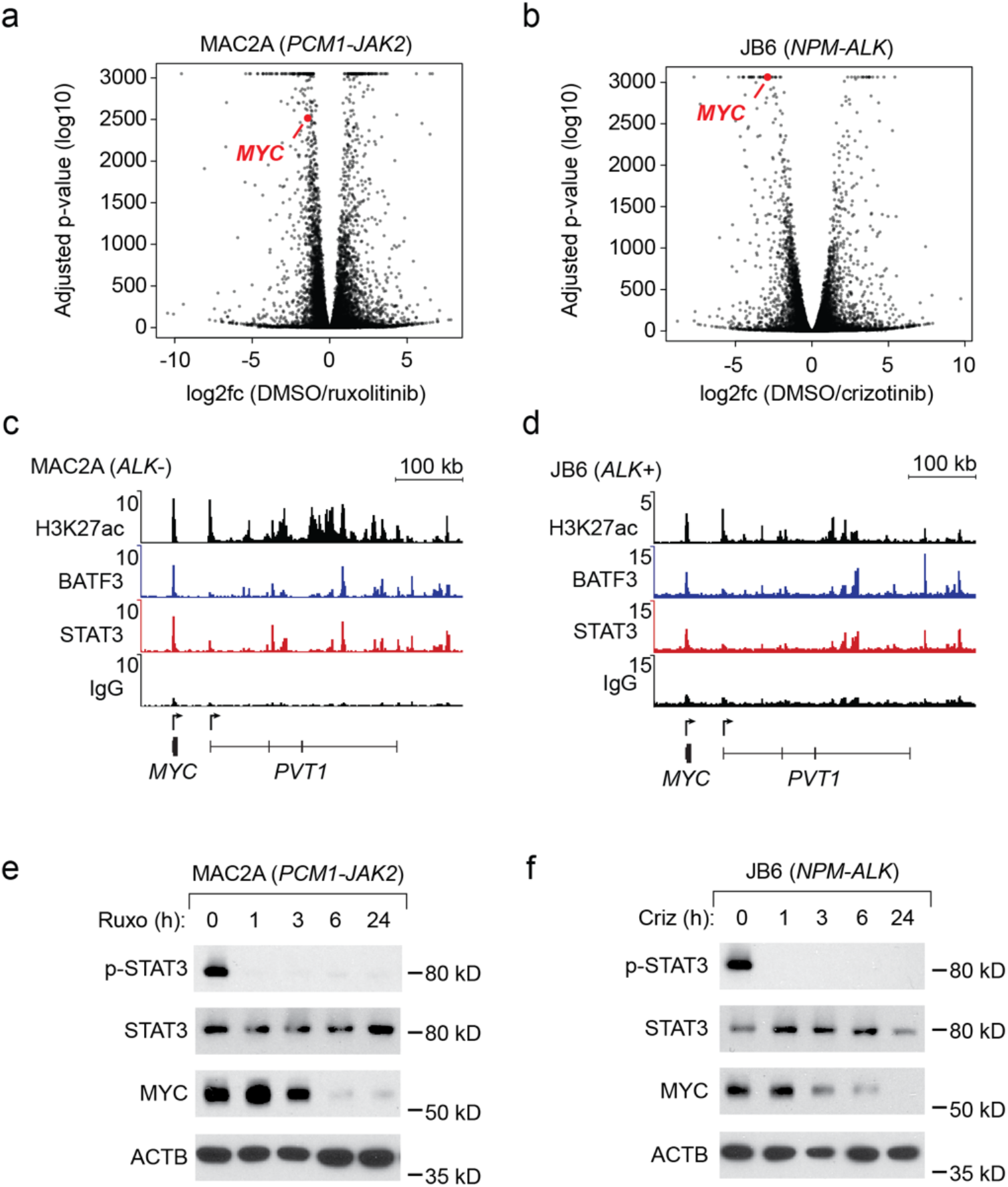
ALK or JAK inhibition attenuates STAT3-driven *MYC* expression. a-b) Volcano plot showing changes in transcript levels when MAC2A cells (a) were treated with ruxolitinib (1 μM, 24 hours) and when JB6 cells (b were treated with crizotinib (1 μM, 24 hours). c-d) ChIP-seq for H3K27ac and CUT&RUN sequencing alignment tracks for BATF3, STAT3 and IgG (control) shown at the *MYC* gene locus in MAC2A (c) and JB6 (d) cells. e-f) Western blot time course showing the protein levels of phospho-STAT3 (Y705), total STAT3, MYC, and ACTB in MAC2A cells (e) treated with ruxolitinib (1 μM) and JB6 cells (f) treated with crizotinib (1 μM) for up to 24 hours.

An immunoblotting time course of MAC2A cells treated with ruxolitinib, and JB6 cells treated with crizotinib, indicated a complete loss of phospho-STAT3 (Y705) by 1 h after treatment initiation (Fig. 4e,f). Furthermore, after the loss of detectable STAT3 phosphorylation, MYC protein levels were significantly reduced, consistent with the essential role of activated STAT3 for high levels of expression of the *MYC* oncogene in ALCL (Fig. 4e,f).

### STAT3 activation is necessary and sufficient for *MYC* expression and ALCL cell survival

To conclusively determine whether activation of STAT3 by NPM-ALK is the critical event required for high levels of *MYC* expression and cell viability in ALCL cells, we tested whether a mutationally activated STAT3 protein could rescue the effects of ALK inhibition in JB6 cells. These cells were transduced with a lentivirus encoding an mEGFP-tagged STAT3^Y640F^-mutant cDNA. Mutation of the *STAT3* gene, such as the gain-of-function Y640F substitution, where the codon for tyrosine-640 is altered to encode for phenylalanine, is often observed as a somatic mutation in ALK-negative ALCL tumors^12^. First, we compared the LD_50_ values of JB6 control (parental) and STAT3^Y640F^-expressing cells when challenged with ALK inhibitors. We found that the LD_50_ is 3-10 fold higher in STAT3^Y640F^ cells for both crizotinib and alectinib, indicating that activation of STAT3 is sufficient to rescue ALCL cell viability downstream of NPM-ALK inhibition (Fig. 5a). Control and STAT3^Y640F^-expressing JB6 cells were then treated with crizotinib and alectinib at 100 nM each, and cell growth was monitored daily for 72 hours. During treatment, control cells rapidly lost viability and failed to proliferate (Fig. 5b). By contrast, STAT3^Y640F^-expressing cells continued proliferating, further demonstrating that direct activation of STAT3 can rescue cell growth and survival of ALK+ ALCL cells treated with an ALK tyrosine kinase inhibitor (Fig. 5b).

**Figure 5.**
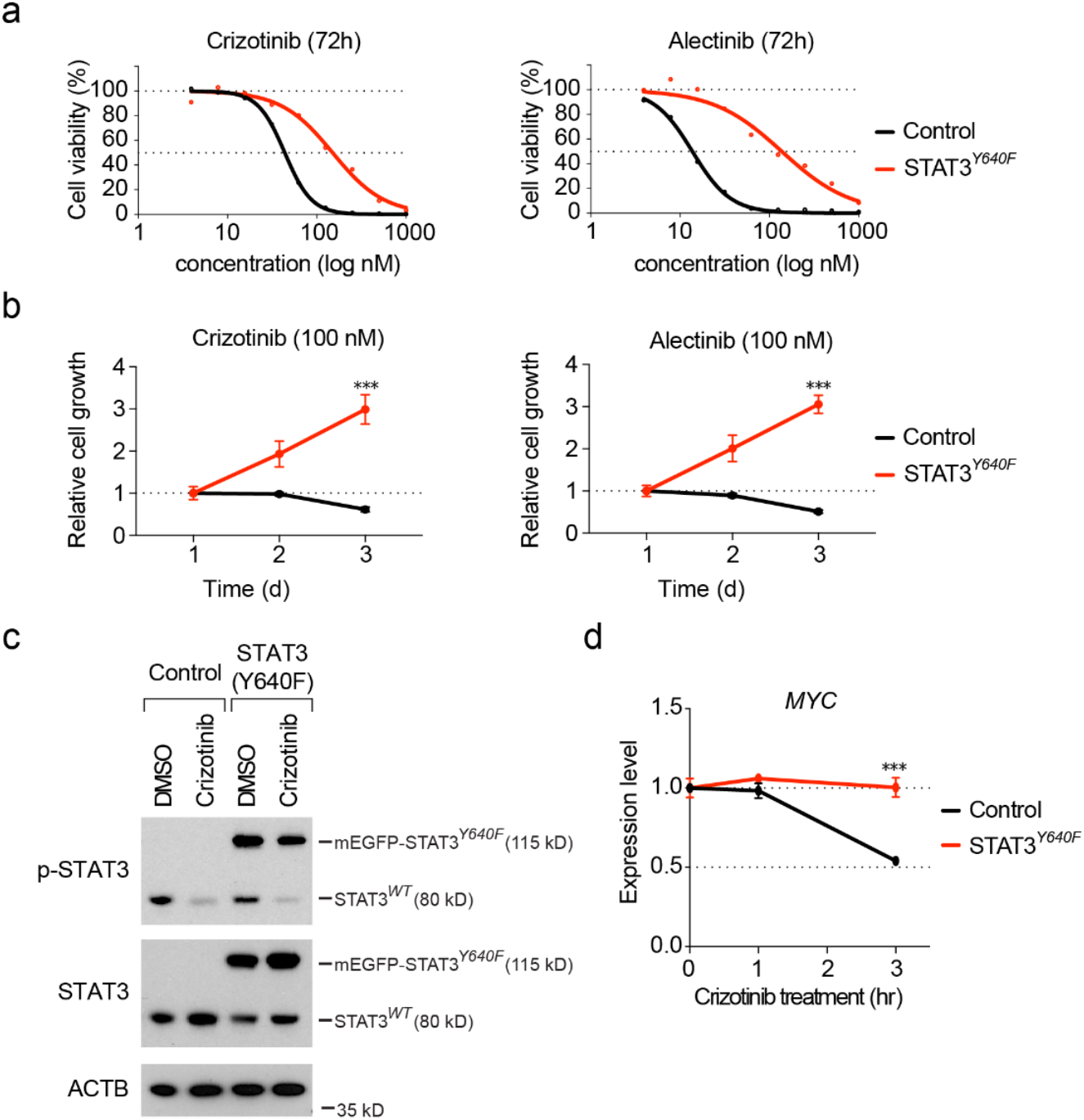
Expression of STAT3^Y640F^ rescues loss of cell growth and viability induced by ALK inhibition. a) Cell viability assays showing relative cell survival of control and STAT3^Y640F^ expressing JB6 cells treated with increasing concentrations of ALK inhibitors – crizotinib or alectinib – for 72 hours. b) Cell proliferation time course assay showing the relative growth of control and STAT3^Y640F^ JB6 cells treated with a single concentration of crizotinib or alectinib (100 nM) for up to 72 hours. c) Western blot for phospho-STAT3 (Y705), total STAT3 and β-actin in control, and STAT3^Y640F^ JB6 cells treated with DMSO, crizotinib (100 nM) for 6 hours. d) Quantitative PCR time course assaying *MYC* gene expression levels in JB6 control and STAT3^Y640F^ cells treated with crizotinib (100 nM) for 0, 1 and 3 hours, demonstrating that STAT3^Y640F^ expression rescued MYC expression at 3 hours (p>0.001).

Next, we sought to assess the biochemical effects of the crizotinib ALK kinase inhibitor on STAT3 phosphorylation in control JB6 cells and cells expressing the STAT3^Y640F^ mutant protein by immunoblotting with a phospho-STAT3 (Y705) antibody (Fig. 5c). Six hours after treatment with 100 nM crizotinib, both the control and transduced JB6 cells exhibited loss of phosphorylation of the endogenous (unmutated) STAT3 protein. By contrast, the STAT3^Y640F^ mutant protein showed no reduction of phosphorylation at Y705 following treatment with crizotinib, indicating that it remains active despite inhibition of NPM-ALK (Fig. 5c). Since our results in Figure 4 indicated that STAT3 is an essential regulator of *MYC* gene transcription, we assayed *MYC* transcript levels by quantitative RT-PCR immediately following treatment with crizotinib in control and STAT3^Y640F^ cells. By 3 h after the addition of 100 nM crizotinib, *MYC* mRNA levels were reduced by 50% in control cells compared to cells expressing STAT3^Y640F^, which retained high levels of *MYC* RNA expression that were equivalent to the levels in untreated cells (Fig. 5d). These results demonstrate that rescuing STAT3 activity with a gain-of-function mutation, resulting in constitutive STAT3 phosphorylation independent of ALK, is necessary and sufficient to retain high levels of *MYC* expression and preservation of ALCL cell viability despite the inhibition of ALK with small molecule kinase inhibitors.

### STAT3 and MYC invade SEs driving CRC transcription factors

MYC family proteins have an essential role in promoting transcription initiation and elongation, and overexpression of MYC in tumor cells causes amplification of the entire transcriptome of the cell^39,40^. Highly abundant MYC proteins in tumor cells have the potential to invade the SEs of highly expressed genes by binding to lower affinity E-boxes, such that MYC increases the transcriptional output and reinforces oncogenic expression programs^41,42^. To assess the genomic co-occupancy of enhancers and SEs by MYC in ALCL, we performed CUT&RUN sequencing for MYC in MAC2A (*PCM1-JAK2*+) and JB6 (*NPM-ALK*+) cells. Along with BATF3 and STAT3, MYC binding was strongly enriched at SEs, including those associated with the CRC transcription factor genes *BATF3, IRF4* and *IKZF1* (Fig. 6a-c), as well as the typical enhancer and promoter driving *STAT3* and large SE region associated with *MYC* (Fig. 6d,e). This indicates that MYC expression also participates in a positive feedback loop, which includes the *STAT3* enhancer and promoter, and functions as a component of the CRC by binding and occupying the SEs that control the expression of itself and each of the other CRC transcription factors (Fig. 6f).

**Figure 6.**
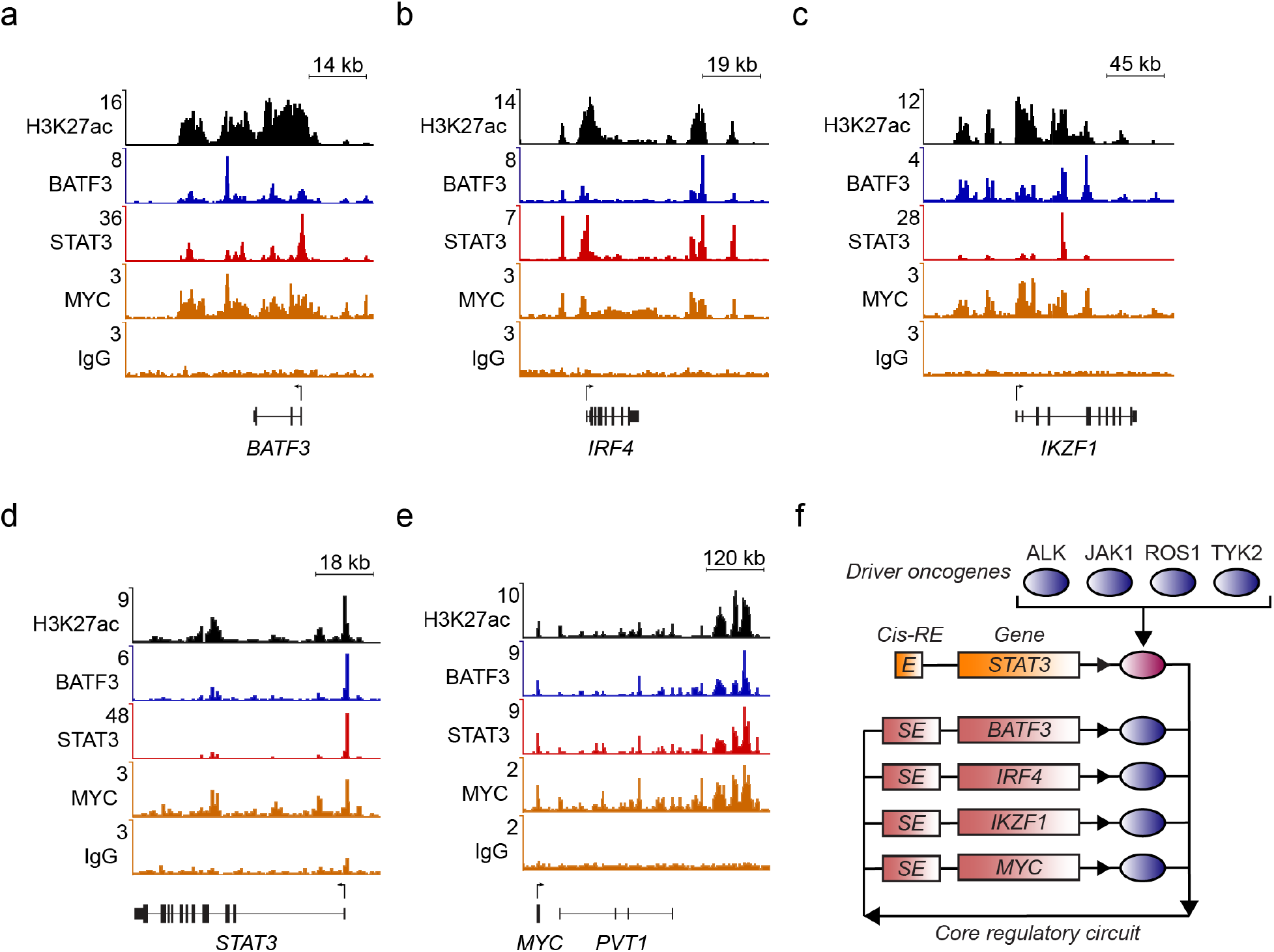
MYC invades SEs driving CRC transcription factors. a-e) H3K27ac ChIP-seq and CUT&RUN sequencing alignment tracks for BATF3, STAT3, MYC and IgG (control) in MAC2A cells, shown at the gene loci for *BATF3* (a), *IRF4* (b), *IKZF1* (c), STAT3 (d) and *MYC* (e). Read densities (y-axis) were normalized to reads per million reads sequenced in each sample. f) Illustration showing how BATF3, IRF4, IKZF1 and MYC form an interconnected co-regulatory loop together with STAT3, which is activated by dysregulation of ALK, JAK, ROS1, or TYK2. Rectangles illustrate regulatory elements and gene loci and oval symbols illustrate proteins.

## Discussion

Aberrant activation of signaling pathways downstream of tyrosine kinase proteins can drive increased cell growth and survival, leading to oncogenic transformation^37,43–46^. In ALCL, previous studies have converged on the JAK-STAT pathway as the central driver of ALCL tumorigenesis, with STAT3 as a critical terminal effector^12^. In NPM-ALK+ ALCL, STAT3 is activated by NPM-ALK either directly – or possibly through intermediate JAK pathway kinases – and is required for cell survival and maintaining the neoplastic phenotype of ALCL cells^47,48^. The ALK-negative subtype of ALCL frequently harbors activating mutations in *JAK1, ROS1, TYK2* or *STAT3* directly, indicating a critical role for STAT3 activation in both subtypes^12^. Although STAT3 activation provides essential pro-growth signals required for ALCL tumorigenesis, it is not fully understood how exactly STAT3 facilitates the malignant identity of ALCL cells.

In this study, we profiled the enhancer landscape of NPM-ALK+ and ALK-negative ALCL by H3K27ac ChIP-seq and identify a conserved set of SEs associated with highly expressed genes that encode transcription factors, several of which have been implicated in the pathogenesis of ALCL^27,49– 51^ (Fig. 1). Among these highly expressed transcription factor genes, we demonstrate that *BATF3, IRF4*, and *IKZF1*, are genetic dependencies in ALCL and act as part of an interconnected autoregulatory loop referred to as the transcriptional core regulatory circuitry (Fig. 2). Core regulatory transcription factors have emerged as highly selective gene dependencies across a broad range of tumor types^18,22,23,52^. The gene regulatory activity of the CRC is essential for driving a lineage-specific transcription program and maintaining cell state^18^. ALCL cells were not only selectively dependent on this small group of core regulatory transcription factors, but they are also dependent on *STAT3* (Fig. 3). Interestingly, STAT3 does not meet our criteria for a CRC component, since its expression is not driven by an autoregulatory SE, rather, it is activated as an independent event through tyrosine phosphorylation. Once activated, STAT3 binds enhancers already occupied by the CRC, which explains why activation of conserved signaling pathways can have disparate effects in different cell types that express unique combinations of CRC transcription factors. Thus, STAT3 belongs to a class of proteins referred to as “signaling transcription factors,” which are activated by signaling cascades and then can bind cooperatively to enhancers to exert conditional effects on the cellular transcriptome^53,54^. In this sense, signaling transcription factors provide an additional level of regulation, allowing cytokine signaling to regulate transcription via enhancer occupied by the CRC. The ability of cytokine binding to launch signaling pathways downstream of cell receptors, like tyrosine phosphorylation of STAT3, may function as an “on-off switch” for activation of the underlying CRC^24^. In the case of ALCL, the receptor signaling aspect usually required for STAT3 activation is subverted by the expression of constitutively active tyrosine kinases that have become disconnected from their normal conditional activation by cytokines produced from surrounding signaling cells.

Here we demonstrate that, once activated, STAT3 binds to the regulatory SEs of the CRC transcription factors *BATF3, IRF4*, and *IKZF1*, as well as the *MYC* SE, driving high levels of *MYC* expression and permitting MYC to co-regulate the ALCL CRC (Fig. 4). *MYC* expression is a critical downstream target regulated in response to the activation of STAT3, which represents a novel mechanism through which ALCL cells activate *MYC* expression. Other tumor types frequently employ direct amplification of the *MYC* gene^55,56^, or enhancer hijacking from highly expressed genes by genomic structural rearrangements^57,58^, ALCL tumor cells appear to employ activation of STAT3 as a mechanism of *MYC* activation. Here we show that the expression of a hyperactive STAT3^Y640F^ mutant protein could rescue *MYC* expression, along with the loss of cell growth and viability observed following treatment with ALK inhibitors (Fig. 5). Unlike the normal STAT3 protein that loses phosphorylation of STAT3 during ALK inhibitor treatment, the STAT3^Y640F^ mutant protein remains phosphorylated at Y705 without ALK signaling. Mechanisms of resistance development against ALK inhibitors remain to be unknown, however, some studies have found that loss of either *PTPN1* or *PTPN2* induces resistance to small molecule ALK inhibitors *in vitro* and *in vivo*, and in ALK inhibitor–relapsed patient tumors, elevated *IL10RA* expression rewires STAT3 signaling even without phosphorylation by NPM1-ALK^34,48^. The essential role of phosphorylated (active) STAT3 in the transformation of ALCL, whether NPM-ALK+ or ALK-negative, suggests that treatments targeting STAT3 directly could be an effective treatment strategy for ALK-negative or ALK-inhibitor resistant ALCL^37,47^.

Activated STAT3 and the transcription factors CRC, form a group of strong tumor-selective gene dependencies in ALCL, which exists independent of *ALK* status, and could form the basis of a safe and effective therapeutic approach for ALCL. Activated STAT3 represents an attractive target because of its requirement for post-transcriptional modification by phosphorylation before it can perform its oncogenic role. As the relationship between oncogenic signaling pathways and transcriptional regulatory proteins becomes further resolved, therapeutic strategies that exploit the interconnectedness of these dependencies are becoming a feasible approach for developing new targeted therapies.

## Acknowledgments

This work was supported by NIH grants R35CA210064 (A.T.L.), R35CA210030 (K.S.) and K08CA245251 (A.D.D), and the St. Jude Children’s Research Hospital Collaborative Research Consortium on Chromatin Regulation in Pediatric Cancer. N.P. was supported by a grant from the Lymphoma Research Foundation. A.D.D. and B.J.A. are supported by the American Lebanese Syrian Associated Charities (ALSAC). M.W.Z. was supported by grants from the Alex’s Lemonade Stand Foundation, Charles A. King Trust, and Claudia Adams Barr Foundation.

## Materials and methods

### Cell culture

ALCL cell lines Karpas299, SUDHL1, JB6, SR-786, SUP-M2, FE-PD, MAC1, and MAC2A were obtained from ATCC. All cell lines were cultured in RPMI-1640 supplemented with 10% FBS and 100 IU/ml penicillin. Cells were tested for mycoplasma every 3 months with the Mycoalert kit (Promega), and identity was confirmed by STR profiling (DFCI molecular diagnostics laboratory). Cell proliferation assays were performed by seeding 5000 cells per well in 96-well plates containing DMSO, crizotinib, or alectinib (MedChemExpress, LLC). Cell viability was assayed with CellTiter-Glo according to the manufacturer’s protocol using a Spectramax M5 plate reader (Promega).

### Antibodies

For CUT&RUN, we used the following antibodies from Cell Signaling: BRD4 (E2A7X), IKZF1 (D6N9Y), IRF4 (D6P5H), STAT3 (D3Z2G), MYC (5605); R&D systems: BATF3 (AF7437); Bethyl Laboratories: MED1 (A300-793A); and Santa Cruz Biotechnology: Mouse IgG (sc-2025). For ChIP-seq, we used the following antibodies from Abcam H3K27ac (ab4729); Cell Signaling: STAT3 (124H6); and R&D systems BATF3 (AF7437). For Western blotting, we used the following antibodies from Cell Signaling: p-STAT3(Y705) (9145), STAT3 (12640), MYC (5605), ACTB (5970).

### Patient-derived xenograft model

Both male and female NSG mice were purchased from the National Cancer Institute (Frederick, MD). Human ALCL patient-derived xenograft cells (line: WCTL-81162-Q13), obtained from the Dana-Farber Center for Patient-Derived Models, were subcutaneously implanted into the hind flanks of nine- to ten-week-old mice. When tumor volumes reached approximately 2000 mm^3^, mice were euthanized, tumor cells were extracted and rinsed in PBS, fixed in PBS containing 1% formaldehyde for 10 min, washed with PBS, and snap-frozen in liquid nitrogen for ChIP-seq experiments.

### Western blotting

Protein samples were collected and lysed using a radioimmunoprecipitation assay buffer containing protease and phosphatase inhibitors (Cell Signaling Technology). Lysates were quantified by Bradford assay (Bio-Rad), and 10 μg of extracted protein was separated using Novex SDS– polyacrylamide gel electrophoresis reagents and transferred to nitrocellulose membranes (Life Technologies). Membranes were blocked in 5% milk protein and incubated with primary antibodies overnight, followed by secondary horseradish peroxidase–linked goat anti-rabbit and anti-mouse (Cell Signaling Technology) antibodies (1:1000) according to the manufacturers’ instructions. Antibody-bound membranes were incubated with SuperSignal West Pico chemiluminescent substrate (Thermo Fisher Scientific) and developed using HyBlot CL autoradiography film (Thomas Scientific).

### Quantitative RT-PCR

Total RNA was harvested using the RNeasy kit (QIAgen) according to the manufacturer’s protocol. First-strand synthesis was performed with Superscript III (Invitrogen).

Quantitative PCR analysis was conducted on the ViiA7 system (Life Technologies) with SYBR Green PCR Master Mix (Roche) using validated primers specific to each target each gene. We used the following primer sequences MYC-F (AGCGACTCTGAGGAGGAACAA), MYC-R (TCCAGACTCTGACCTTTTGCC), ACTB-F (AGAGCTACGAGCTGCCTGAC), and ACTB-R (AGCACTGTGTTGGCGTACAG).

### RNA-seq

RNA isolation was performed using the RNeasy Mini Kit (Qiagen), and RNA quality was assessed on a Fragment Analyzer (Advanced Analytical Technologies) − SS Total RNA 15nt. RNA-seq libraries were prepared using the Kapa mRNA HyperPrep Kit for Illumina (Roche) with Poly(A) selection according to the manufacturer’s instructions. Library quantification was examined on a Fragment Analyzer − HS NGS Fragment 1-6000bp and Qubit HS dsDNA Kit (Invitrogen). Libraries were pooled and sequenced to 150bp paired-end on the Illumina NovaSeq platform. Sequencing data were analyzed as described previously^59^.

### RNA-Seq analysis

Raw reads were aligned to the hg19 revision of the human reference genome to which the sequences of the ERCC spike-in probes were added using hisat2^60^ v2.1.0 in paired-end mode. Expression was quantified for all RefSeq genes downloaded 5/17/17 using htseq-count with parameters -i gene_id --stranded=reverse -m intersection-strict. Differential expression was determined statistically using DEseq2 and read counts from each sample individually^60^.

### ChIP-sequencing

ChIP-seq was performed as previously described^50^. For each ChIP, 5 μg of antibody coupled to 2 μg of magnetic Dynabeads (Life Technologies) was added to 3 ml of sonicated nuclear extract from formaldehyde-fixed cells. Chromatin was immunoprecipitated overnight, cross-links were reversed, and DNA was purified by precipitation with phenol:chloroform:isoamyl alcohol. DNA pellets were resuspended in 25 μl of TE buffer. Illumina sequencing, library construction, and ChIP-seq analysis methods were previously described. Reads were aligned to the human reference genome (hg19) using bowtie v1.2.2 with parameters –k 2 –m 2 –best and –l set to the read length. For visualization, WIG files were created from aligned read positions using MACS v1.4 with parameters –w –S –space=50 –nomodel – shiftsize=200 to artificially extend reads to 200 bp and to calculate their density in 50-bp bins. Read counts in 50-bp bins were normalized to the millions of mapped reads, giving RPM values.

### ChIP-Seq and CUT&RUN-Seq processing

For patient-derived xenograft models, reads were first aligned to the mm9 revision of the mouse reference genome using bowtie v1.2.2 in paired-end mode with parameters -k 2 -m 2 – best and the non-mapping reads retained using –un. For remaining reads in the xenograft samples and all reads for other samples, reads were aligned to the human reference genome (hg19) using bowtie v1.2.2^61^ with parameters –k 2 –m 2 –best and –l set to the read length. Paired-end samples were aligned in paired-end mode. For visualization, WIG files were created from aligned read positions of single-end read samples using MACS v1.4^62^ with parameters –w –S –space=50 – nomodel –shiftsize=200 to artificially extend reads to 200 bp and to calculate their density in 50-bp bins. Read counts in 50-bp bins were normalized to the millions of mapped reads, giving RPM values. WIG files were visualized in the IGV browser version 2.7.2. For coverage tracks, paired-end alignments were converted into fragments using samtools^63^ view, samtools sort, and bedtools^64^ bamtobed. Fragment coverage in 50-bp bins was calculated by building a genome-wide list of bins using bedtools makewindows and bedtools intersect. Per-million-fragment signal was created by dividing by the number of mapped fragments.

### Super-enhancer identification and assignment

Super-enhancers in ALCL cells were identified in each cell line separately as previously described^65^ using ROSE (https://bitbucket.org/young_computation/rose). Reads overlapping ENCODE-defined ignorable regions were removed from both H3K27ac and input BAM files using bedtools intersect against ENCFF001TDO. From the remaining reads, two sets of peaks of H3K27ac were identified using MACS and the positions of aligned reads with parameter sets –keep-dup=auto –p 1e-9 and –keep-dup=all –p 1e-9. The collapsed union of regions called using these MACS parameter sets were used as input for ROSE with parameters - s 12500 -t 1000 -g hg19. Enhancers were assigned to the single expressed gene, defined as being in the top two-thirds of the promoter (TSS ± 500 bp) H3K27ac coverage in a sample, whose transcription start site (TSS) was nearest the center of the enhancer.

### Coverage heatmaps

Heatmaps showing coverage of purified sequence fragments (i.e. ChIP-Seq and CUT&RUN-Seq) were built at 4kb windows centered on the collapsed union of peaks for transcription factors. Peaks were identified using MACS v1.4 with corresponding control and reads and parameters -p 1e-9 – keep-dup=auto. Coverage of reads was quantified in these regions using bamToGFF (https://github.com/BradnerLab/pipeline/blob/master/bamToGFF.py) with parameter -m 50. Rows (regions) were ordered by the row sum of H3K27ac signal.

### CRISPR-Cas9 dependency screen analysis

Public data from genome-scale CRISPR-Cas9 screens performed at the Broad Institute were downloaded from FigShare (https://figshare.com/articles/dataset/DepMap_22Q2_Public/19700056/2). A total of n=1086 cancer cell lines (including n=5 ALCL lines) were screened with the Avana library, containing 73,372 guides and an average of 4 guides per gene^66^. The screens were conducted in a pooled experiment as previously described^29,67^. Genetic dependencies that are enriched in ALCL cell lines were identified using linear-model analysis from the limma v3.38.3 R package^68^ by performing a two-tailed t test for the difference in distribution of gene dependency scores in ALCL compared to all other cell lines screened as previously described^28^. Statistical significance was calculated as a q value derived from the p value corrected for multiple hypothesis testing using the Benjamini & Hochberg method^69^. Transcription factor genes were highlighted based on a published list of human transcription factors^70^.

## Data and Code Availability

Raw and processed data files were deposited to the NCBI GEO server under super-series GSE158916 and GSE212077. Code written in R/python to perform analyses of ChIP-seq and CUT&RUN is available upon request.

## Additional statistics

Data from the ChIP-seq experiments were analyzed as described above. Cell viability data were analyzed with two-way ANOVA followed by post-hoc t-test. Statistical significance was defined as a p-value <0.05. Data were analyzed with GraphPad Prism 9.4.0, and all error bars represent SD unless otherwise noted.

